# Low-dose ethanol produces mixed effects on cortico-hippocampal theta rhythms and modifies hippocampal theta frequency through multiple mechanisms

**DOI:** 10.1101/2019.12.25.888487

**Authors:** Calvin K. Young, Neil McNaughton

**Author notes:** **Corresponding Author:** Dr. Calvin K. Young, Department of Psychology, University of Otago, PO BOX 56, Dunedin, New Zealand.

## Abstract

Ethanol is one of the most widely used drugs – with many psychoactive effects, including anxiolysis. The deleterious effects on brain function and general health of chronic and high-level ethanol use are well-studied. However, the neurophysiology of acute low dose ethanol has not been systematically investigated. Here, we examined the effects of low dose (0.25 and 0.5 g/kg) ethanol on midline (prefrontal, cingulate and retrosplenial) cortical and hippocampal theta oscillations in freely moving rats. We also tested low dose ethanol on reticular-elicited and running-elicited hippocampal theta frequency and assessed the linear relationship of theta frequency to stimulation intensity and running speed, respectively. Low dose ethanol had mixed effects on cortical theta oscillations. It most reliably reduced theta frequency, produced a weak inverted-U effect on theta power, and had no detectable effect on cortico-hippocampal theta coherence. Ethanol dose-dependently decreased the y-intercept of the speed-theta frequency function without affecting the slope, but decreased the slope of the stimulation intensity-theta frequency function without affecting the y-intercept; thus decreasing theta frequency in both cases but through different mechanisms. We conclude low dose ethanol has weak but detectable effects on cortical and hippocampal theta oscillations. These effects may underlie positive cognitive and behavioural outcomes reported in the literature using low dose ethanol. The double dissociation of slope and y-intercept specific changes relating to different methods of hippocampal theta elicitation suggests that multiple mechanisms contribute to anxiolytic effects on theta and so hippocampal function.

## Introduction

Alcohol is one of the most widely used drugs. Frequently, humans use ethanol as self-medication for symptoms of anxiety (Buckner et al., 2008, Robinson et al., 2009); and there are related effects with its consumption by rodents (Blanchard et al., 1993). However, for clinical use, ethanol’s deleterious effects and abuse potential far outweigh its efficacy as an anxiolytic; and its effects on defensive behaviour are more complex than benzodiazepines such as diazepam (Blanchard et al., 1993).

How ethanol’s many actions on the brain interact to produce physiological and behavioural changes is still poorly understood (Abrahao et al., 2017, Cui and Koob, 2017, Tizabi et al., 2018). While the deleterious health consequences of chronic, heavy use of alcohol are well-established, the impacts of low-dose ethanol on brain physiology and on behavioural outcomes are less clear (Cui and Koob, 2017, Tizabi et al., 2018).

Early work implied a biphasic relationship between plasma ethanol and behaviour. Variation in the delivered dose, and the resulting blood alcohol levels across time, produced results that suggest that low levels tend to have “activating” effects, while high levels are “suppressive” (Pohorecky, 1977). Human studies have shown that ethanol has negative effects on mnemonic (potentially hippocampal) and executive (potentially frontal) functions at doses as low as 0.5 g/kg (Dry et al., 2012, Day et al., 2015). But, in rodents, low doses (<0.75 g/kg) of ethanol can increase locomotion (Phillips and Shen, 1996, Karlsson and Roman, 2016), facilitate fear conditioning (Gulick and Gould, 2007), and improve spatial working memory (Rossetti et al., 2002). Interestingly, theta oscillations in the HPC and in various parts of the cortex (especially the prefrontal cortex; PFC) appear to be functionally linked during locomotion (Young and McNaughton, 2009), fear conditioning (Lesting et al., 2011), and spatial working memory (Jones and Wilson, 2005, Benchenane et al., 2010). However, it is not known how an improvement of function is related to neural changes elicited by low dose ethanol. A single previous report, recording from an unspecified *frontal* location, found no effect on brain oscillations with up to 0.5 g/kg (Prado de Carvalho and Izquierdo, 1977). In contrast, a modest increase in *hippocampal* theta power and frequency has been reported at 0.25 and 0.5 g/kg (Givens, 1995).

As with other types of anxiolytics (e.g. benzodiazepines, buspirone), ethanol has been shown to reduce the frequency of reticular (RPO)-elicited hippocampal (HPC) theta oscillations (5-12 Hz) in rodents (Coop et al., 1990, McNaughton et al., 2007). Ethanol reduces the slope of the linear relationship between RPO stimulation intensity and HPC theta frequency likely via an action on a benzodiazepine site (Coop et al., 1990). The RPO-elicitation paradigm is highly robust and specific in terms of predicting human clinical anxiolytic action (McNaughton and Sedgwick, 1978, Gray and McNaughton, 2000, McNaughton et al., 2007), and has been used in a range of laboratories as a screen of known or potential anxiolytic drugs (Engin et al., 2009, Siok et al., 2009, Siok et al., 2012, Yeung et al., 2012).

More recently, anxiolytics have also been shown to reduce HPC theta frequency related to the speed of locomotion (Wells et al., 2013) – a well-established and robust behavioural correlate of HPC theta oscillations (Vanderwolf, 1969, McFarland et al., 1975, Slawinska and Kasicki, 1998, Robbe et al., 2006, Hinman et al., 2011, Wells et al., 2013). Despite this superficial similarity in effects of anxiolytics, the RPO-elicitation paradigm explicitly requires an absence of overt motor movements (Gray and McNaughton, 2000), and hence is likely to depend on somewhat different circuits than movement-elicited theta (Kramis et al., 1975, Sainsbury et al., 1987).

Differences also exist in how each anxiolytic reduces HPC theta frequency. In the locomotion paradigm, anxiolytics appear to reduce HPC theta frequency solely by decreasing the y-intercept of the speed-frequency relationship without affecting the slope (Wells et al., 2013); an effect largely replicable in the entorhinal cortex (Monaghan et al., 2017). In contrast, in the RPO paradigm, anxiolytics may decrease the y-intercept and/or slope of the stimulation intensity-frequency relationship (McNaughton and Sedgwick, 1978, Gray and McNaughton, 2000, McNaughton et al., 2007). It is currently not understood how anxiolytics reduce RPO- and running-elicited HPC theta frequency, given that the two methods appear to differ in the underlying mechanisms of their generation (Kramis et al., 1975, Bland, 1986).

The effects of low doses of ethanol have not been studied on hippocampal theta elicited by any means nor on the interactions between hippocampus and cortex on which its cognitive and behavioural effects must depend. Here, we tested in freely behaving rats the impact of low (0.25 and 0.5 g/kg) ethanol on: 1) hippocampal versus cortical theta oscillations; and 2) hippocampal theta frequency elicited by RPO stimulation intensity versus by speed of locomotion.

## Methods

### General procedures

Male Sprague-Dawley rats were obtained from the Otago University Animal Breeding Station. The rats were group-housed prior to surgery and testing. Up to 14 days were given for acclimatization and handling. Water and food were available ad libitum and the rats were kept on a 12/12 hour light/dark cycle (lights on at 6 am) at 21+/-2° C. Post-surgery, rats were housed singly under the same conditions. Ethical approval was obtained from the Otago University Animal Ethics Committee (approval number 35/04).

Two types of electrodes were used in the current study. Twisted bipolar electrodes were made from Teflon-coated stainless steel wires (200 μm in diameter, A-M Systems, Inc., USA). Multi-array electrodes were made from 100 μm Teflon-coated stainless steel wires, bonded by using Loctite 401 (Loctite Australia Pty. Ltd., Australia). A detailed description can be found in Young and McNaughton (2009). All electrodes were tested for electrical continuity and impedance before interfacing to a McIntyre mini-connector.

The animals were anaesthetised with sodium pentobarbital (60 mg/kg) or with ketamine and medetomidine (0.75 mg/kg and 0.5 mg/kg, s.c.). Betadine (Faulding Pharmaceuticals, Australia) was applied to the skin and Tricin (Jurox Pty. Ltd., Australia) applied to the eyes. Electrodes/arrays were inserted into the brain (coordinates and surgical procedures are given below) and dental cement was used to hold and anchor the electrodes and arrays into place. Sutures were applied to the wound after the dental cement application. If ketamine/medetomidine was used for anaesthesia, 0.5 mg/kg of atipamezole was given as an antidote to medetomidine to facilitate recovery. All animals were given at least twelve days of recovery before testing.

At the end of the experiment, the animals were euthanised with halothane, and transcardially perfused with 0.9% saline solution followed by 10% formalin in 0.9% saline. The brain was removed and kept in 10% formalin in dH_2_O overnight and then fixed with 30% sucrose in 10% formalin. Upon sucrose saturation, the brains were sliced (90 μm) on a freezing microtome, mounted on gelatine-coated slides, stained with thionin, covered with DPX mounting solution, and cover-slipped. Histological reconstruction was done under a microscope or a microfiche reader. Array reconstruction relied on the determination of the deepest point of the implant and known tip separation within the array, supported by high-resolution photographs of the arrays prior to implant.

### Open field cortico-hippocampal recordings

Rats (n = 19) received bipolar HPC and midline neocortical array electrode implants. Table 1 gives target coordinates; exact placements are reported in Young and McNaughton (2009). Ethanol administration was counterbalanced by splitting the animals into a group of 9 and a group of 10. Ten percent ethanol in a 0.9% saline solution was given in ascending order at doses of 0.0 g/kg (saline), 0.25 g/kg and 0.5 g/kg *i.p.* to one group and in descending order to the other group. Inter-session interval was >48 hours to allow wash-out. Out of 19 rats, one did not have a correct HPC placement, one had noisy recordings not suited for further analysis, and one had a missing video. Data from these three rats were excluded and sixteen rats contributed to the analysed data reported here.

**Table 1.** Neocortical implant coordinates for all rats included in the analysis. All distances (AP; anterior-posterior, ML; medial-lateral, DV, dorsal-ventral) are in mm and referenced against bregma. Note two arrays were in the RS for CY54.

| Subject | Six-Electrode Array |  |  |  | Four-Electrode Array |  |  |  |
| --- | --- | --- | --- | --- | --- | --- | --- | --- |
|  | AP | ML | DV | Angle | AP | ML | DV | Angle |
| CY16 | 1.70 | 1.50 | 3.60 | 15 | - | - | - | - |
| CY27 | 1.70 | 1.50 | 3.60 | 15 | -4.16 | 1.50 | 2.20 | 25 |
| CY29 | 1.00 | 1.50 | 3.00 | 15 | -4.80 | 1.50 | 2.20 | 25 |
| CY30 | 0.20 | 1.40 | 2.70 | 15 | -5.80 | 1.50 | 3.00 | 0 |
| CY33 | 2.20 | 2.00 | 5.00 | 16 | 2.30 | 1.40 | 2.60 | 21 |
| CY34 | -0.80 | 1.40 | 2.70 | 19 | -6.80 | 1.50 | 2.60 | 20 |
| CY35 | 1.70 | 1.40 | 3.40 | 16 | -3.30 | 1.40 | 2.20 | 21 |
| CY36 | 1.00 | 1.40 | 3.10 | 16 | -5.20 | 1.40 | 2.10 | 22 |
| CY41 | 3.20 | 2.00 | 5.40 | 16 | -4.16 | 2.40 | 2.30 | 21 |
| CY44 | 1.70 | 2.40 | 3.40 | 17 | -4.80 | 2.40 | 2.20 | 23 |
| CY45 | 1.70 | 1.70 | 3.60 | 6 | -5.20 | 2.40 | 2.10 | 22 |
| CY46 | 2.20 | 2.30 | 5.00 | 0 | -2.30 | 2.40 | 2.60 | 21 |
| CY47 | 1.00 | 1.70 | 3.30 | 6 | -3.30 | 2.40 | 2.20 | 21 |
| CY48 | 0.20 | 1.70 | 2.90 | 7 | -5.80 | 2.50 | 3.00 | 21 |
| CY49 | -0.80 | 2.40 | 2.70 | 19 | -6.80 | 2.50 | 2.60 | 20 |
| CY50 | -1.30 | 1.40 | 2.40 | 20 | -8.00 | 1.50 | 2.50 | 8 |
| CY51 | 1.70 | 1.40 | 3.40 | 16 | -6.30 | 1.50 | 2.70 | 21 |
| CY53 | 2.20 | 2.00 | 5.00 | 16 | -1.80 | 2.00 | 3.00 | 27 |
| CY54 | -7.64 | 1.50 | 3.50 | 8 | -1.50 | 2.00 | 3.00 | 27 |

Cortical and HPC LFPs were fed through an impedance transformer and then amplified by an EEG-4214 unit (Nihon Kohden, Japan). Data were then digitized via a National Instruments (Austin, TX) DAQ system (PCI-MIO-16E-4 interfaced through SCB-68) and sampled at 128 Hz using a custom-written LabVIEW programme. Behavioural testing apparatus and procedures for array-implanted rats have been described previously (Young and McNaughton, 2009). Briefly, rats were connected to the recording system and placed in a 73 x 63 x 51 cm box with black interior walls. Each rat received four 6-minute exposures/trials in a single session.

Position tracking (25 frames/s) was carried out offline using manually corrected trajectories generated by iTracker (Perez-Escudero et al., 2014). The trajectories were processed further by linear interpolation, followed by a five-point median filter. Distance, and hence speed, was calculated by approximating pixels-to-centimetres based on known open field dimensions over time. We considered speeds over 2.5 cm/s as locomotion and used 2.5 cm/s speed bins to account for the differences in speed distributions between trials (Wells et al., 2013). In sum, speeds between 2.5 to 20 cm/s (inclusive of 90% of locomotion data) in 2.5 cm/s bins were used for further analysis.

Multi-taper spectral analyses were carried out using the Chronux (RRID:SCR_005547) package (Bokil et al., 2010), with three tapers and a bandwidth product of five. All power, frequency, and coherence measures were derived from 5-12 Hz from the spectra. Instantaneous frequency was calculated by first deriving the analytical signal from a Hilbert transform of band-passed (5-12 Hz) LFPs, smoothed by using a 3rd order Savitzky-Golay filter, before taking the derivative as the estimation of the instantaneous frequency. As an estimation of dominant frequency, a weighted mean of theta power (centre of mass; CoM) was calculated. A least-squares linear fit was also used to fit the speed/theta frequency relationship, with slope, intercept and estimated theta frequency (as derived from fitted speed value between 2.5 and 20 cm/s) were used for further analyses. Linear least-squares fit was applied to theta power/CoM to locomotion speed to examine their inter-dependencies.

### RPO stimulation

Rats (n = 6) received bipolar recording electrodes implanted in the dorsal subiculum (AP: −6.0, ML: 2.0, DV: −3.5 mm; 0.5 mm tip separation) and bipolar stimulating electrodes targeting the output fibres of the RPO (AP: −7.0, ML: 1.6, DV: −8.0 mm; 0.5 mm tip separation). The recording electrodes were connected to a field-effect transistor (FET) impedance transformer that was connected to an amplifier (Grass P511K, USA). The amplifier was connected to a Hitachi Storage Oscilloscope V-134 (Hitachi, Japan) and a National Instrument interface card (NI BNC 2110, USA) connected to a PC via a PCI-MIO-16E-4 data acquisition device (National Instruments, USA). The stimulating electrodes were operated by a Grass SD9 stimulator which was also connected to the BNC 2110 and PCI-MIO-16E-4 interface cards. This setup allowed the stimulation and recording to be controlled via LabVIEW (National Instruments, USA). Rats were placed in a cylindrical enclosure (circumference = 100 cm, height = 22 cm). A custom program was written in LabVIEW to trigger stimulation and record the data. Stimuli were delivered as a 100 Hz, 0.1 ms monophasic square pulse for 0.8 s at various voltages. The program captured 0.5 s of the post-stimulation LFP and had a custom-made peak finding algorithm for wave counting. The first three (theta) cycles were used to determine frequency based on peak-to-peak intervals.

To observe the relationship between theta frequency and stimulation intensity, four or five equally-spaced intensity steps were used for each animal (ranging from 0.5-1.5 V). The number of steps and the level of increment were dependent on the range of stimulation intensity of each rat, but none exceeded 10 V or produced any stimulus-bound motor activity (e.g. locomotion or unilateral head/body tilt). Once this stimulation range was established, the rats were tested within this range in both ascending and descending orders to ascertain a baseline. This baseline was used to verify that elicited hippocampal theta frequency was linearly proportional to the intensity of RPO stimulation. Rats that: 1) failed to exhibit HPC theta upon RPO stimulation; 2) displayed motor activity upon theta elicitation; 3) did not yield changes in theta frequency that were dependent on stimulation intensity; 4) had a stimulation range that was too small (<2 V); or 5) yielded >7.5 Hz theta LFP at the lowest stimulation intensity were not tested further. Two out of six rats were excluded in this fashion. Full dose range data were obtained and analysed from four rats. Four additional rats tested against 0.0, 0.25 and 0.5 g/kg were used for analyses involving those doses only.

The rats were given 0.0, 0.25, 0.5, 1 or 2 g/kg ethanol intraperitoneally in 0.9% saline (with the ethanol concentration at 10% or 20% depending on the volume needed to achieve the dose) twenty minutes after the first trial that served as the baseline. Five more trials followed at post-injection times of 10, 40, 70, 100, and 130 minutes. For each trial, the animals were tested with their individual stimulation intensity range in ascending order then in descending order. This was repeated twice more in the same fashion to produce six sets of replicated measurements per trial. After all trials were completed, the rats were returned to their cages and were given at least a 48 hour wash-out period before the next testing session.

A least-squares linear fit was applied to assess the relationship between stimulation intensity and theta frequency. Slope and intercept from the fit were extracted for further analysis. RPO-elicited theta frequency was then taken as the frequency intercept of the median intensity from the testing range of RPO stimulation intensities.

### Statistical analyses

All data were submitted to either mixed-model ANOVA with polynomial contrasts or Pearson’s correlation analysis. All *p* values reported are uncorrected unless otherwise specified.

## Results

### Cortico-hippocampal theta oscillations in the open field

Firstly, we assessed the spectral profile of hippocampal and cortical recording sites. Figure 1A illustrates the distribution of recording sites and the number of rats (n =16) contributing to each recording site. Figure 1B shows the spectral profile (2 - 20 Hz) when the rats’ movement speed was > 2.5 cm/s (red) or < 2.5 cm/s (black). Theta power and dominant frequency both appear to be higher for periods when the movement speed is > 2.5 cm/s. In the HPC, there is a clear difference in the power (locomotion, F(1,94) = 9.42, *p* = .003) and dominant theta frequency (locomotion, F(1,94) = 23.86, *p* < .001) of theta oscillations between nominal movement/no movement (CA1; Figure 1B). For the most anterior recording areas (PFC; Figure 1B), there may be marginal increases in theta power (n.s.) but no obvious theta frequency shifts (n.s.) during periods of locomotion. However, despite lack of statistical difference in theta power between locomotion states, there was an increase of theta coherence with the hippocampus (PFCxCA1; Figure 1B; locomotion, F(1,64) = 8.03, *p* = .006). More caudally, there is little evidence to suggest a difference in theta power (n.s.) or frequency (n.s.) in the Cg (Cg; Figure 1B), nor is there a significant change in theta coherence between the Cg and CA1 (CgxCA1; Figure 1B). The RS showed clear theta oscillations during locomotion (RS; Figure 1B) with an increase in dominant theta frequency between nominal movement/ no movement (locomotion, F(1,101) = 7.03, *p* = .009) but no reliable increase in power (locomotion F(1,101) = .831, p = .364, n.s.). The frequency increase was also accompanied by an increase in theta coherence with the HPC (RSxCA1; Figure 1B; locomotion, F(1,101) = 6.92, *p* =. 01).

**Figure 1.**
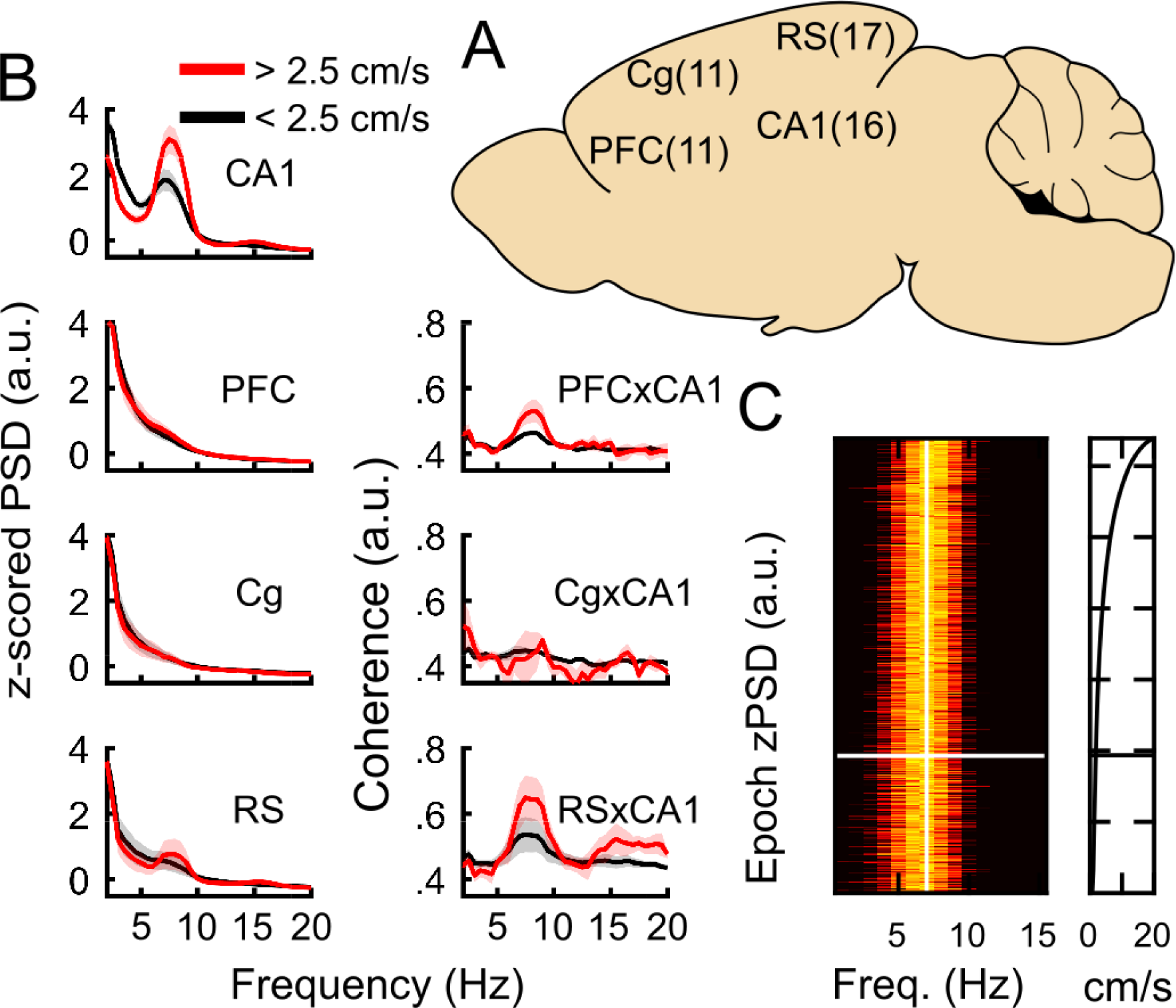
Movement modulation of theta oscillations in the cortex. (A) Number of recording sites across cortical parcellation adopted in the current study. (B) Changes in 0-20 Hz activity across the cortices between no movement (black, < 2.5 cm/s) and locomotion (red > 2.5 cm/s). Data displayed as mean (solid lines) and standard error (shaded area). (C) Speed-ordered, normalised power spectral density (left) with a vertical white line aligned to 7 Hz and a horizontal white line aligning the 2.5 cm/s cut off. The distribution of speeds included in the analysis (right; 2.5 to 20 cm/s) matching speed-ordered PSDs on the right. CA1, cornu ammonis area 1; PFC, prefrontal cortex; Cg; cingulate cortex; RS, retrosplenial cortex; PSD, power spectral density.

After establishing theta oscillations can be observed in our re-derived bipolar cortical recordings, we sought to investigate the relationship between movement speed and theta oscillations in the HPC and cortical recording sites. We observed a correlation between spectral profiles of HPC LFPs between 0-15 Hz as a function of movement speed (Figure 1C), but a linear relationship is only evident at speeds over 2 cm/s (grey line). Particularly, we found that HPC instantaneous theta frequency in the 5-12 Hz range increased as a function of movement speed between 2.5 to 20 cm/s (r(17660) = .19, *p* <.0001). In the rest of the cortices only PFC (r(8089) =. 07, *p* <.0001) and RS (r(5253) =. 08, *p* <.0001) show weak (<1% of the variance) but statistically significant locomotion speed modulation of instantaneous theta frequency, but not the Cg (r(6375) = .003, *p* = .40). There was virtually no theta power modulation by locomotion speed (highest r =. 08 across all regions), including HPC (r(17660) =. 04, *p* <. 0001).

### Behavioural and neurophysiological effects of low dose ethanol

Behaviourally, EtOH appeared to dose-dependently decrease mean locomotion speed as testing session progressed (dose [lin] x time [lin], F(1, 12) = 10.44, *p* <. 01; Figure 2A). In contrast, saline control did not result in changes in the amount of locomotion over time, 0.25 g/kg EtOH increased the amount of locomotion as the session progressed and 0.5 g/kg resulted a decrease (dose [quad] x time [lin], F(1, 12) = 6.74, *p* <. 05; Figure 2B).

**Figure 2.**
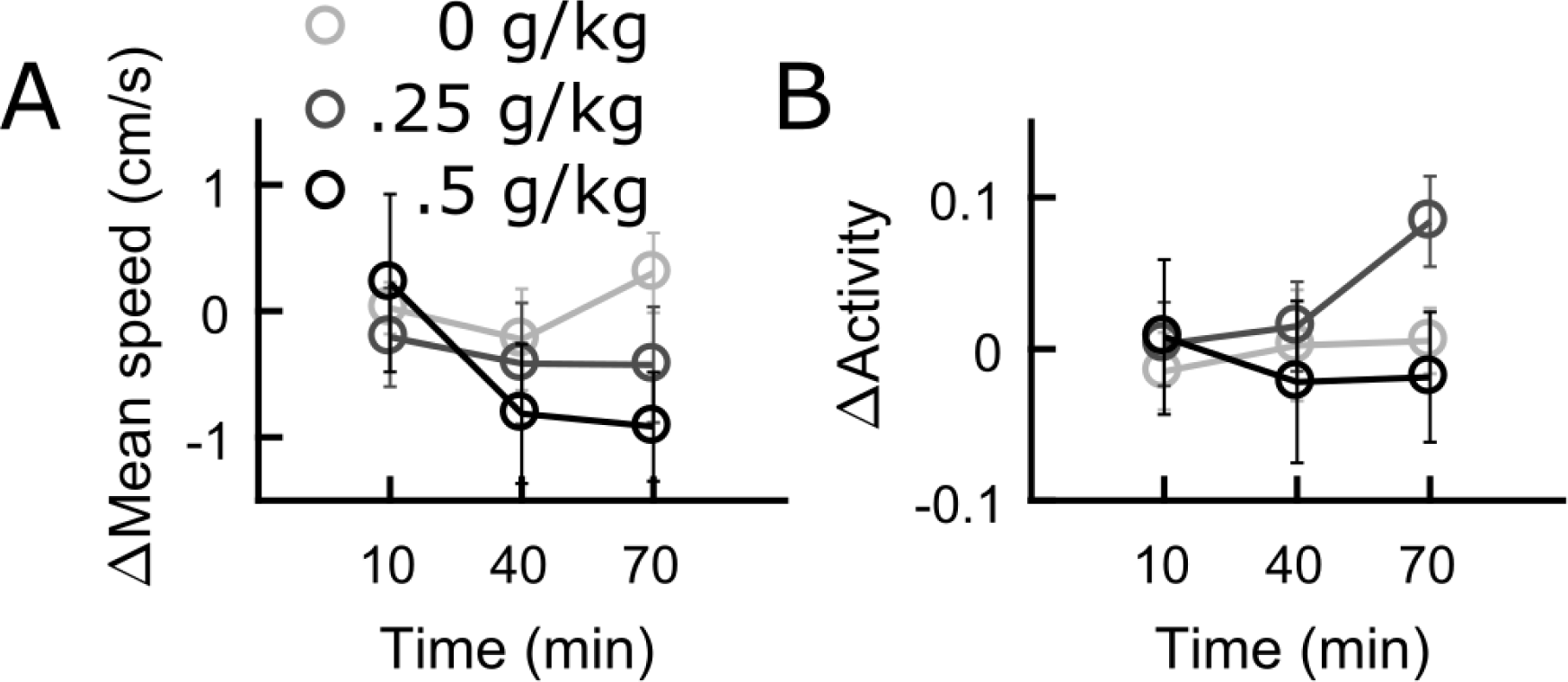
Locomotion measures in the open field in response to low dose ethanol. (A) change in mean speed referenced against the pre-injection session. (B) Change in activity, as measured by the amount of locomotion, in response to ethanol.

Previous studies have shown blood ethanol concentration to peak within the first 20 minutes (Givens, 1995, Rossetti et al., 2002), hence we used our 10 min post-injection time point to assess the effects of EtOH on theta power, frequency and coherence across the recorded cortical regions. EtOH appears induce an inverted-U change in theta power in all cortical areas (Figure 3A). However, only changes in the RS (black line) reached statistical significance (dose [quad], F(1,4) = 6.28, *p* = .03). In contrast, EtOH had a mostly linear effect of dose on theta frequency (Figure 3B), with the exception of the PFC, where a statistically marginal U-shaped trend can be observed (dose [quad], F(1,9) = 4.35, *p* = .07). Particularly, 0.25 and 0.5 g/kg decreased Cg theta frequency (dose [lin], F(1,8) = 7.12, *p* = .03), while the RS (dose [lin], F(1,14) = 17.51, *p* < .001) and HPC (dose [lin], F(1,13) = 9.1, *p* = .01) linear trends were stronger. Lastly, there may have been a simple dose-dependent increase in PFC-HPC theta coherence (dose [lin] F(1,9) = 4.15, *p* = .07), but an apparent U-shaped dose-coherence relation for Cg-HPC was not reliable (dose [quad], F(1,7) = 1.13, *p* =.29). There was no observable change in RS-HPC theta coherence with either of the low doses (dose [lin], F(1,14) = .04, *p* =.83).

**Figure 3.**
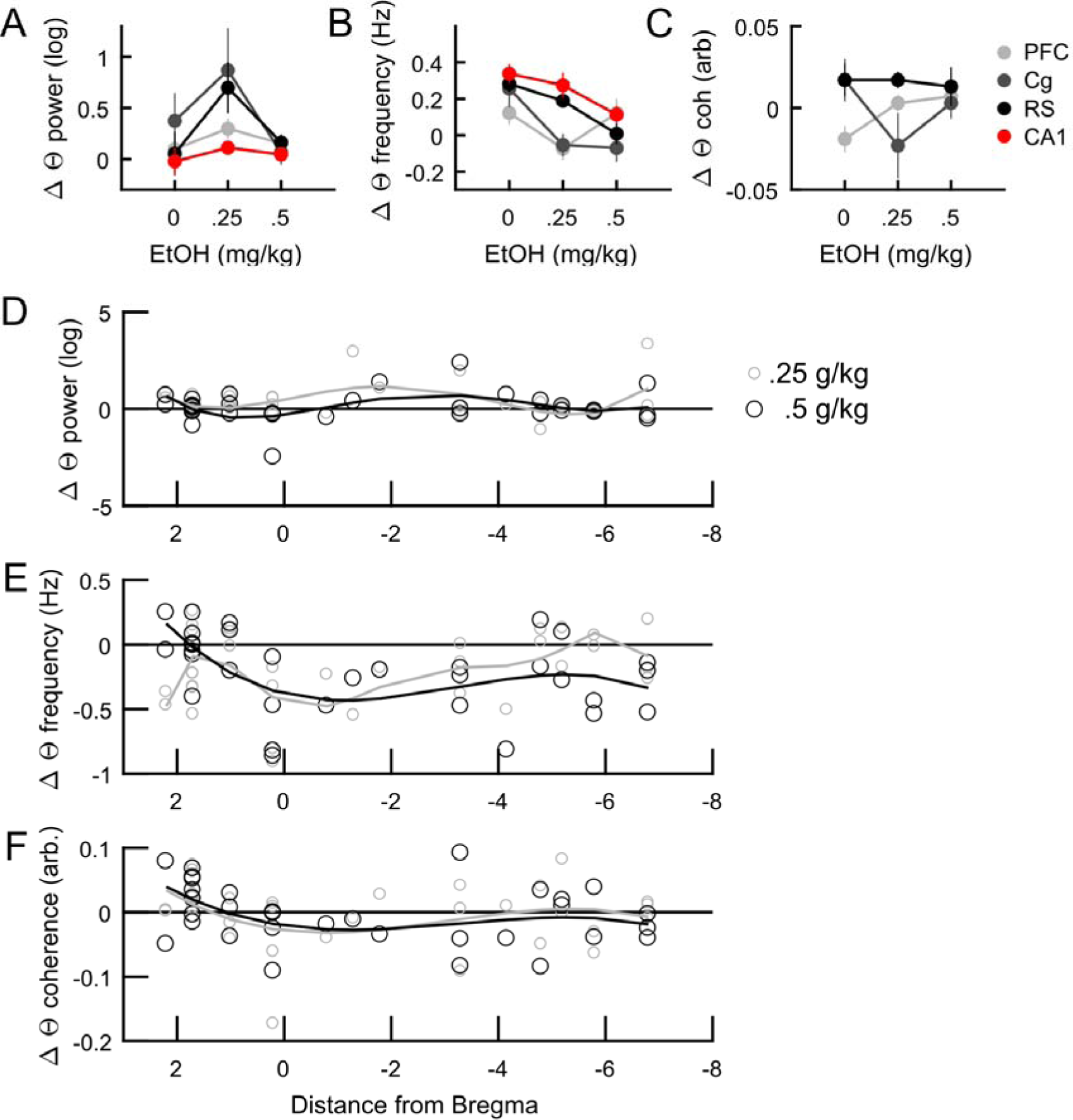
Changes in theta power (between −10 min and +10 min injection), frequency and coherence across different cortical regions in response to low dose ethanol. (A) In all areas, 0.25 g/kg ethanol increase theta power but 0.5 g/kg decreased it. Only changes in RS was found to be statistically significant. (B) 0.25 g/kg reduced theta frequency in all areas. However, 0.5 g/kg decreased CA1 and Cg theta frequency, marginally decrease theta frequency in the RS and did not appear to have an effect on PFC theta frequency. (C) Ethanol also had divergent effects on cortico-hippocampal theta coherence, with no effect on CA1-RS theta coherence, increasing CA1-PFC theta coherence and a biphasic effect n CA1-Cg theta coherence. (D) Change in theta power across the anterior-posterior axis. Both doses are best-fit with 4^th^ order polynomials. (E) Same as in (D) but illustrating changes in theta frequency. 0.25 g/kg was fitted with a 6^th^ order polynomial but largely overlaps the cubic function fitted over 0.5 g/kg. (F) Same as in (D) but illustrating changes in theta coherence. Both doses are fitted with cubic functions. CA1, cornu ammonis area 1; PFC, prefrontal cortex; Cg; cingulate cortex; RS, retrosplenial cortex.

Since our neocortical recording site designation collapses actual recording sites within conventionally defined areas (Paxinos and Watson, 2007), we sought to investigate if any relationships exist across the anterior-posterior axis (the main axis of variation in electrode implant positions). HPC recording sites were not part of this analysis. Best polynomial fits, as determined by the lowest order that markedly reduced residuals for the fit compared to the previous, but offers little improvement compared to the next (data not shown). For theta power (Figure 3D), changes across the AP axis mirror those reported in Figure 3A, where minimal changes are seen in the most anterior cortical sites, with a marked increase through the Cg to the rostral part of the RS for both 0.25 and 0.5 g/kg (best fit by 4^th^ order polynomials). In addition, the AP mapping of neocortical theta power changes suggests the increase of 0.25 g/kg induced increase in Cg and RS probably arise from increases mapped onto −2 to −4 mm from bregma. The overall decrease in neocortical theta frequency is clear in Figure 3E, where most data points fall below the baseline. Again, the AP mapping mirrors findings reported in Figure 3B, where a U-shaped relationship between dose and theta frequency can be seen at the most anterior sites (corresponding to the PFC), followed by consistent decreases across the Cg and RS, with some indication of a dose-dependent effect at AP = −5 to −8. Although the best fit for 0.25 g/kg is of the 6^th^ order polynomial and 0.5 g/kg the 3^rd^ order, clear overlap of the fit lines suggest both doses resulted in the largest theta frequency reduction in the Cg. Again, AP-mapped neocortical-HPC theta coherences (Figure 3F) are consistent with data reported in Figure 3C, where a slight but statistically non-significant increase on PFC-HPC theta coherence is consistent at the most anterior recording sites. However, large variability within the extent of Cg and RS suggest inconsistent effects of low dose EtOH on the regions, rather than a lack of effect. Again, both doses are best fit by largely overlapping cubic functions, suggesting both doses induced similar effects across the AP axis in the midline neocortices.

### Effects of low dose ethanol on RPO-versus locomotion-elicited theta oscillations

In rats implanted with HPC and cortical recording probes, we investigated the effects 0.25 and 0.5 g/kg EtOH on locomotion speed-related HPC theta frequency. Interestingly, repeated exposure during the same day appeared to dramatically increase the y-intercept of the locomotion speed vs theta frequency relationship (Figure 4A, left), which did not appear to be affected by 0.25 g/kg EtOH (Figure 4A, middle). However, despite an elevated baseline, 0.5 g/kg EtOH appeared to attenuate y-intercept increase in subsequent sessions (Figure 4A, right). There is a dose-dependent attenuation of theta frequency increase as a result of repeated testing (Figure 4B; dose [lin] x time [quad], F(1,11) = 29.00, *p* <.001), which was mirrored by intercept changes (Figure 4C; dose [lin] x time [quad], F(1,11) = 25.46, *p* <.001), but not slope (Figure 4D; dose [lin] x time [quad], F(1,11) = .03, *p* = .87).

**Figure 4.**
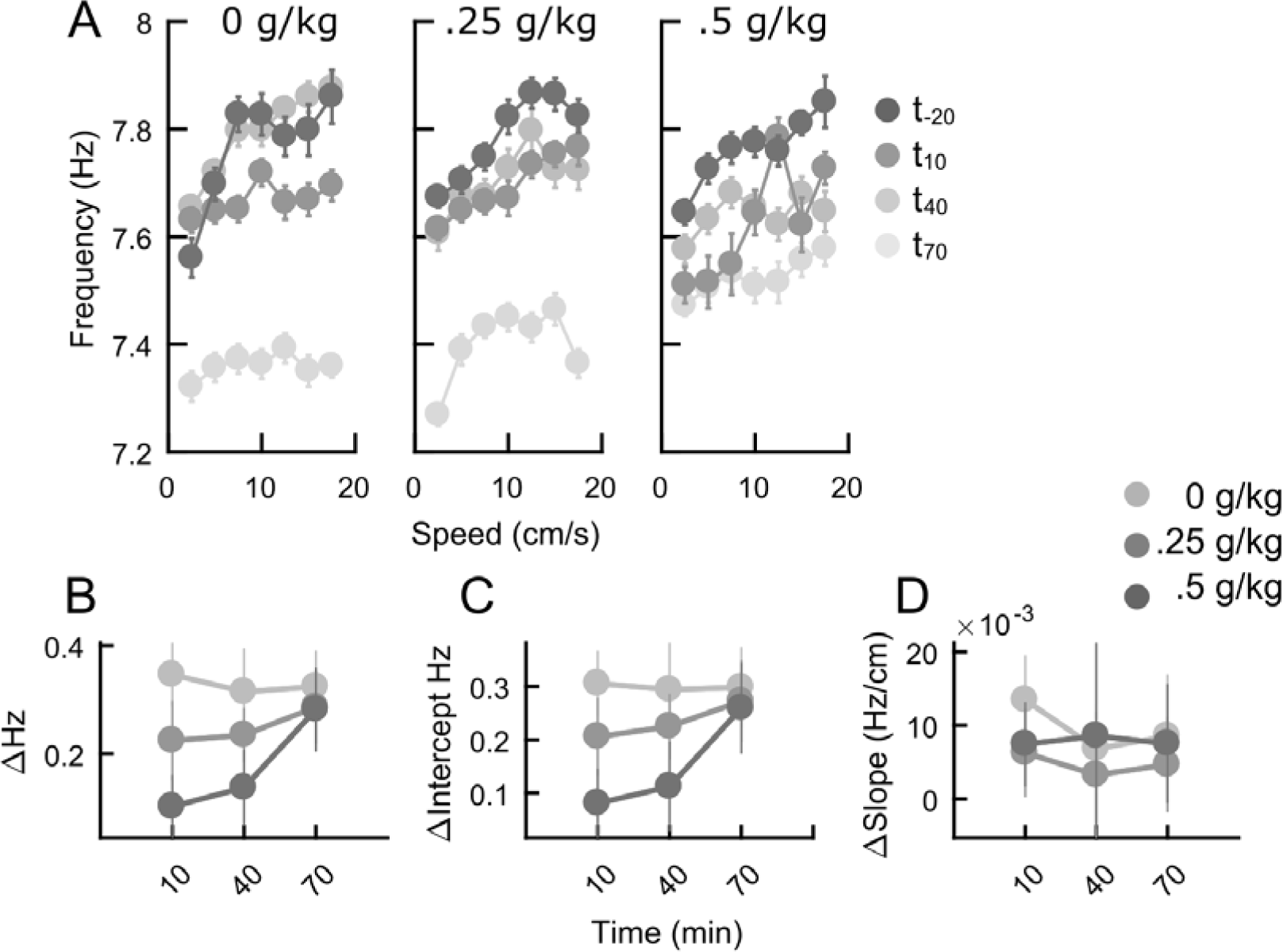
Effects of low dose ethanol on the linear relationship between hippocampal theta frequency and running speed. (A) Population (n = 16) data on repeated exposures to the open field arena (greyscaled data points) with 0.0 g/kg (left), 0.25 g/kg (middle) and 0.5 g/kg (right) injections. (B) Timecourse of mean theta frequency changes, as measured by the mid-point of the linear fit over theta frequency-running speed relationships, in response to low dose ethanol challenge. (C) Timecourse of y-intercept changes of the same linear relationship described in (B). Timecourse of slope changes of the same linear relationship described in (B).

In a separate set of experiments (Figure 5A), we first replicated previous findings on how EtOH affects RPO-elicited HPC theta (Coop et al., 1990), and extended it to include the 0.25 and 0.5 g/kg doses used above. From rats showing a stable relationship between RPO-stimulation intensity and HPC theta frequency (n = 4), we examine the difference between baseline (20 minutes prior to EtOH injection) and post-EtOH (10 minutes after EtOH injection) epochs. There is some indication that 0.25 g/kg increases RPO-elicited HPC theta frequency, consistent with Givens (1995). Recovery towards baseline is only evident for doses lower than 2 g/kg (Figure 5B). The dose-dependent recovery after 130 min was found to be statistically significant (dose [cub] x time [lin], F(1,3) = 100.00, *p* = .002). Theta frequency slope (Figure 5C) was similarly affected (dose [cub] x time [lin], F(1,3) = 12.29, *p* = .04) but intercept was not (Figure 5D).

**Figure 5.**
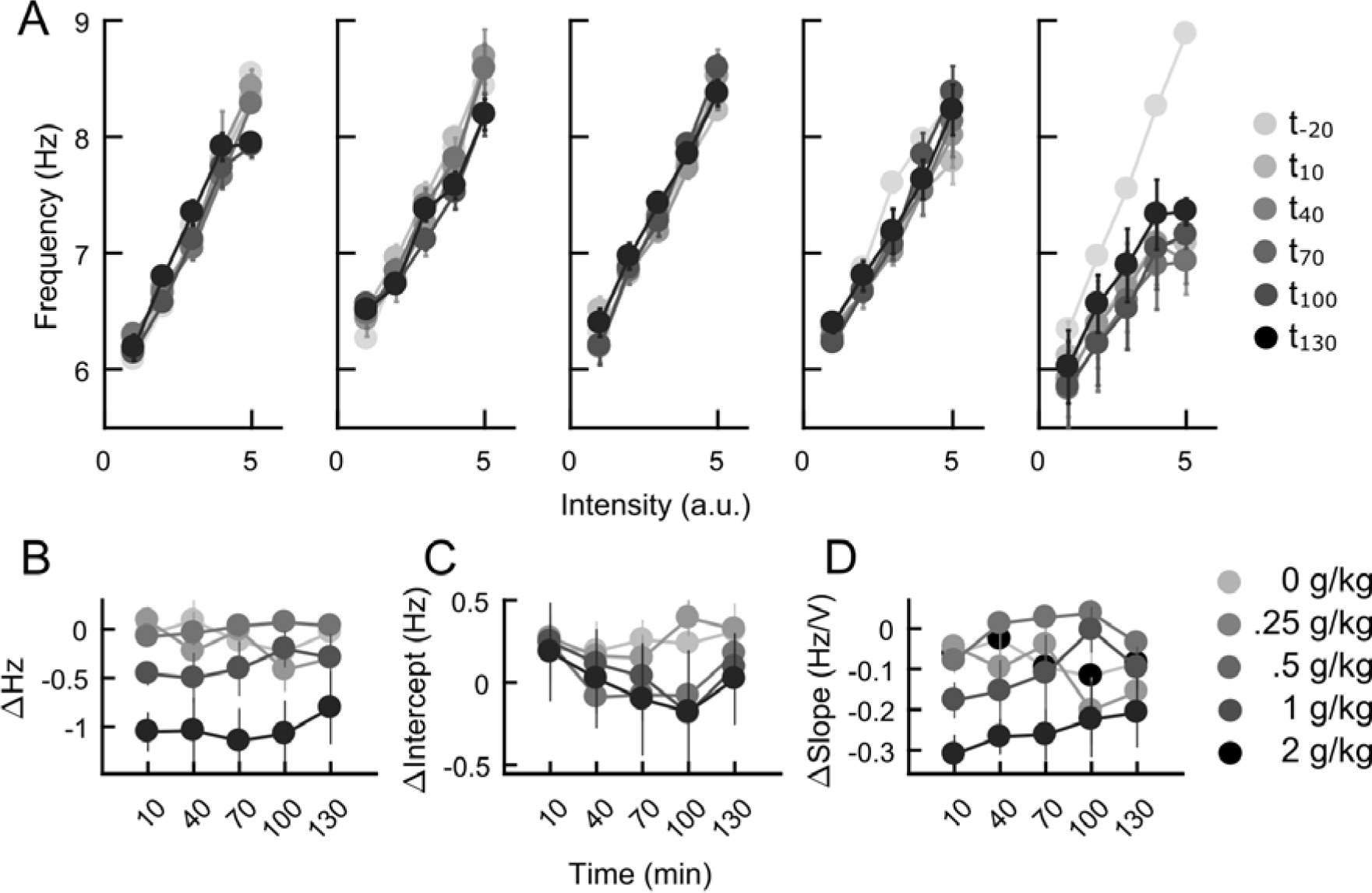
Effects of low dose ethanol on the linear relationship between hippocampal theta frequency and reticulat stimulation intensity. (A) Population (n = 4) data depicting the linear relationship between suprathreshold reticular stimulation intensity and elicited hippocampal theta frequency in increasing ethanol dose (left to right) and at different testing points (greyscale data points). (B) Timecourse of mean theta frequency changes, as measured by the mid-point of the linear fit over theta frequency-reticular stimulation intensity relationships, in response to low dose ethanol challenge. (C) Timecourse of y-intercept changes of the same linear relationship described in (B). Timecourse of slope changes of the same linear relationship described in (B).

Our data indicate that ethanol effects on HPC theta frequency are via slope modification for stimulation intensity (Figure 5), and via intercept modification for speed of locomotion (Figure 4). To further investigate this apparent double-dissociation, we correlated changes in theta frequency with changes in slope and intercept. As expected from data from Figure 5, changes in theta frequency induced by RPO stimulation are not correlated with changes with the intercept of stimulation frequency versus theta frequency (Figure 6A; r(46) = .15, *p* =.39), but are correlated with the slope (Figure 6B; r(46) = .78, *p* < .001). In contrast, consistent with data presented in Figure 4, theta frequency changes as a function of locomotion speed are correlated with the y-intercept (Figure 6C; r(22) = .96, *p* < .001) but not the slope (Figure 6D; r(22) = -.01, *p* = .95).

**Figure 6.**
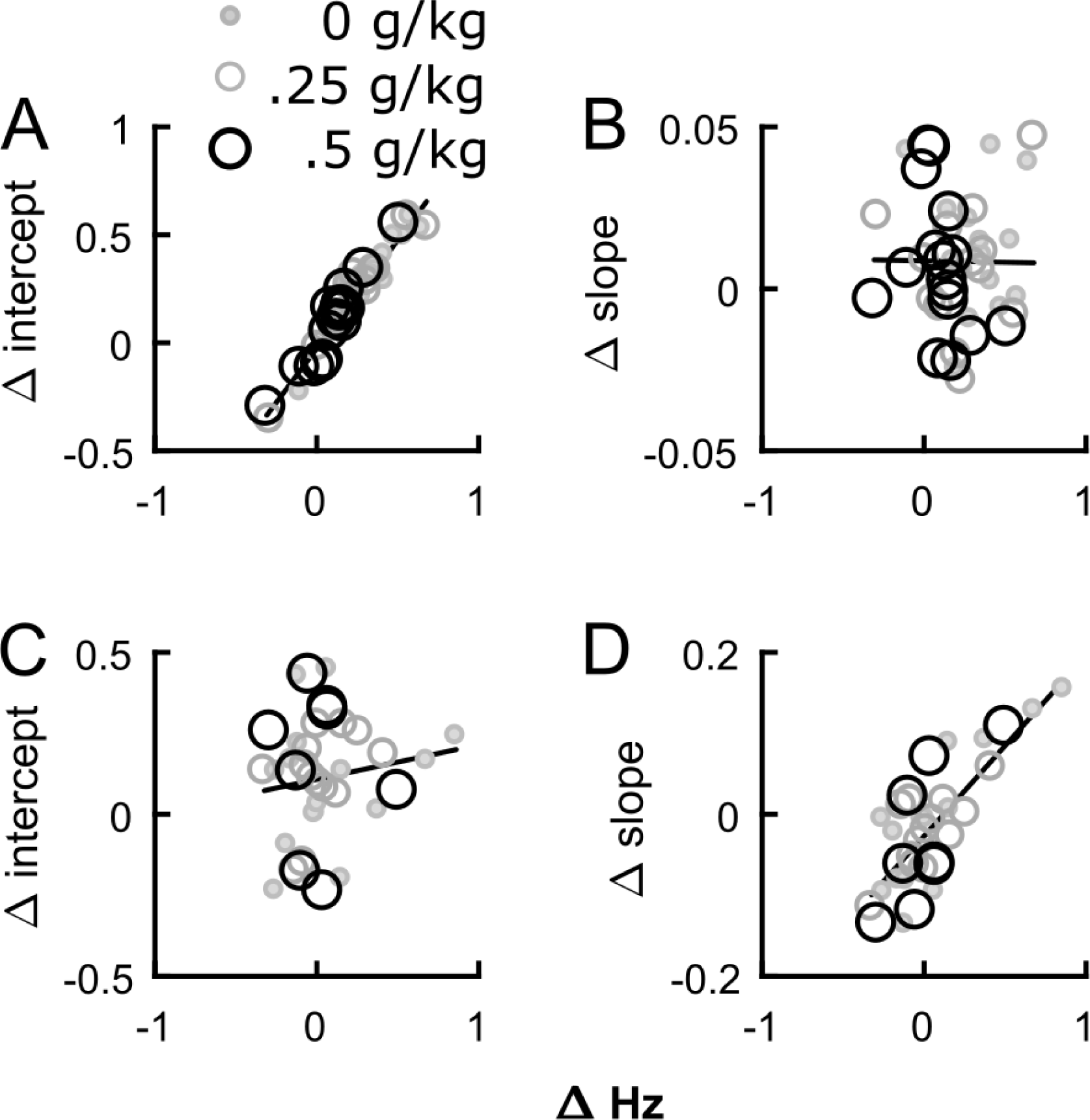
Ethanol modifies hippocampal theta frequency through changing the slope or y-intercept in an elicitation paradigm-dependent manner. (A) Changes in theta frequency are highly correlated with changes in the y-intercept of the running speed vs. theta frequency function (n = 16), (B) but not with slope. (C) Conversely, changes in theta frequency are not correlated with changes in the slope of the reticular stimulation intensity vs. theta frequency function (n = 8), (D) but is highly correlated with the slope of the function instead.

### Discussion

Our main goal was to examine how low dose ethanol: 1) impacts on HPC and cortical theta oscillations and; 2) modifies HPC theta frequency produced by RPO stimulation intensity versus by speed of locomotion. We found modest but consistent biphasic theta power changes: with 0.25 g/kg increasing theta power, whereas EtOH dose monophasically decreased theta frequency throughout the cortex. There were, however, no detectable neocortex-HPC theta coherence changes. There is an indication that Cg is most sensitive to low dose EtOH effects. In addition, we found a weak biphasic effect of low dose EtOH on RPO-elicited theta frequency, but a linear attenuation of theta frequency increase based on locomotion speed. The changes in HPC theta frequency induced by low dose EtOH reveal a double dissociation of how EtOH impacts on HPC theta frequency in the two different elicitation paradigms.

We were able to replicate and extend several fundamental findings reported previously. These address the reliability of source localisation, locomotion-frequency relationships, and effects of ethanol on RPO-elicited theta

The current study differs from our previous report that used a partially overlapping dataset (Young and McNaughton, 2009) in both signal referencing (bipolar re-reference versus monopolar) and cortical parcellation. Nonetheless, we obtained essentially the same pattern of theta distribution across the midline cortices and the same interactions with the HPC. Volume conduction is a contentious issue in EEG/LFP research. It is intrinsic to electrical brain recordings and renders ambiguous the source of the recorded activities. Re-referencing to produce local bipolar signals results in improved representation of local population activities. While local bipolar recording does not completely eliminate volume conduction, we believe our current results suggest that our previous results on theta coherence in un-drugged rats (Young and McNaughton, 2009) indeed reflected midline cortical-HPC interactions and were not dominated by common noise.

With HPC, we were also able to replicate a robust locomotion speed versus theta frequency relationship (McFarland et al., 1975). However, with midline cortex, we found little evidence for a strong relationship. Both PFC and RS theta frequencies were significantly related to locomotion speed, but with extremely shallow slopes and substantial scatter. Currently, the EC is the only area outside the HPC to display a similar speed modulation of local theta LFP frequency (Jeewajee et al., 2008, Hinman et al., 2011, Newman et al., 2013, Monaghan et al., 2017). A lack of general correlation between neocortical theta frequency and locomotion speed is consistent with previous results from the posterior cortex in the rat (McFarland et al., 1975). The lack of speed modulation of midline cortical theta LFPs may be due to a lack of overlap between medial septal sub-regions that project to the HPC-EC and rest of the midline cortices (Gaykema et al., 1990). In fact, there is evidence that RS theta oscillations persist or are sometimes facilitated by medial septal or HPC lesions (Borst et al., 1987, Talk et al., 2004). While PFC theta is thought to be unidirectionally driven by the HPC, only circumstantial evidence exists (O’Neill et al., 2013), and its occurrence can be observed in the absence of HPC theta oscillations (Young and McNaughton, 2009). Overall, we report midline cortical theta to be largely unrelated to locomotion speed and thus functionally distinct from the medial septal-HPC/EC system, which appears to be preferentially involved in spatial context processing.

Lastly, we were able to replicate and extend our knowledge of how EtOH modifies RPO-elicited HPC theta. Our previous study reported that low doses (< 1.7 g/kg) of EtOH did not produce any clear effect on RPO-elicited HPC theta in pilot data involving two rats (Coop et al., 1990). Our current data partially support these findings: clear effects of EtOH on RPO-elicited HPC theta did not appear until 1 g/kg, and were most apparent at 2 g/kg. At these doses, reliable anxiolytic effects can be observed (Gray, 1977, Dalterio et al., 1989). At lower doses, the effects were small with a nonlinear dose-frequency relationship. Particularly, 0.25 g/kg appeared to increase theta frequency whereas 0.5 g/kg produced the same change as saline (0 g/kg), while higher doses linearly decreased theta frequency. These changes match changes in spatial working memory performance using the same doses of EtOH (Givens, 1995). Our function relating dose (i.e. 0.25 – 2 g/kg) to HPC theta frequency also closely resembles that for the ratio of lever presses for food rewards made during a shock-paired tone as a test for anxiety using the same dose range (Dalterio et al., 1989).

Behaviourally, we reported a dose-dependent decrease of movement speed and an inverted-U dose effect on the amount of locomotion over 70 minutes after EtOH/saline administration. However, we did not detect any statistically significant changes in movement speed or the amount of locomotion 10 minutes after EtOH administration, when the effects of low doses of EtOH should be at their peak (Givens, 1995, Rossetti et al., 2002). Likewise, the statistically significant effects detected were related to the changes over time, which appear to outlast the effect of 0.25 and 0.5 g/kg EtOH (Givens, 1995, Rossetti et al., 2002). Our demonstration of a lack of initial effect, and a time-dependent decrease of locomotion in general at 0.5 g/kg EtOH is consistent with previous studies (Chuck et al., 2006, Karlsson and Roman, 2016). The activating or suppressive effects of low dose EtOH on locomotion are less clear in rats, especially between different strains (Criswell et al., 1994, Quintanilla, 1999). Our data indicate no acute effects at presumed peak EtOH concentration, and longer-term (<70 min) trends are likely mediated by secondary causes such as active metabolites of EtOH (Correa et al., 2003).

A major motivation for our study was to address the gap in the literature regarding how low (<1 g/kg) dose ethanol may modify brain oscillations in the cortex. Consequently, there is little prior data for direct comparison. In the HPC, a single previous report with similar low doses to the current study found modest effects with 0.25 and 0.5 g/kg EtOH increasing HPC theta power (Givens, 1995) in an inverted-U function similar to the current study. However, they also described an increasing trend for HPC theta frequency, which is in the opposite direction to our findings. The most obvious difference between the studies is the behavioural context, where Givens employed a spatial working memory paradigm that required locomotion motivated by appetitive rewards, whereas we provided no appetitive rewards or overt cognitive demands. Intuitively, it is most likely that the motor requirements in the spatial working memory paradigm would result in higher locomotion speeds in general and hence increase measured theta frequency, compared to free exploration in the current study.

At higher (>1 g/kg) doses, a general ‘slowing’ of oscillations in the form of increased lower frequency (< 4 Hz) power coupled with a decrease in higher (5-15 Hz) frequency power is often described for the neocortex. For example, in mouse equivalents of Cg and RS in our study, 1.4 g/kg EtOH reduced the power and frequency of oscillations between 0-25 Hz (Ryan et al., 1979). Likewise, 0.8 g/kg in rabbits increased slower (.5-4 Hz) oscillation power at the expense of 5-15 Hz activities in the PFC (Czarnecka and Pietrzak, 1996). Similar but more modest theta power decrease can be observed in the rat in frontal and parietal areas in a dose-dependent (0.5 to 4 g/kg) manner (Young et al., 1982). At 1 g/kg, EtOH may reduce peak frequency in the 4-6 Hz band recorded from frontal and parietal cortices while increasing spectral power only in frontal sites (Slawecki, 2002); however 0.5 g/kg did not result in any observable changes in the frontal cortex (Prado de Carvalho and Izquierdo, 1977). Our data extend the effects of EtOH described in past studies in behaving animals on reducing theta-ranged oscillations in the cortex to 0.5 and 0.25 g/kg, with the exception of PFC, where 0.5 g/kg appeared to increase theta frequency. The biphasic change in theta power is largely due to the increase at 0.25 g/kg; power changes were minimal compared to 0.0 g/kg. This is somewhat consistent with past studies finding no change at 0.5 g/kg (Prado de Carvalho and Izquierdo, 1977, Young et al., 1982). We also report large individual variability in theta modification, particularly in the Cg, in response to EtOH. This variability may have contributed to the apparently contradictory findings on low dose ethanol in previous studies. The marginal increase in PFC-HPC theta coherence reported here is of some interest in the context of the facilitation by low dose EtOH in behavioural tasks where PFC-HPC theta synchrony appears to be involved (Jones and Wilson, 2005, Young and McNaughton, 2009, Benchenane et al., 2010, Lesting et al., 2011)

Our attempt to map the action of low dose EtOH on HPC theta frequency resulted in an interesting dissociation in how the two different theta-eliciting paradigms are affected. A biphasic trend of HPC theta frequency change in the RPO stimulation paradigm was found to be linked to slope changes in relation to stimulation intensity. An attenuation of HPC theta frequency increase during locomotion is associated with the y-intercept of locomotion speed. The apparent larger effects of low dose EtOH on movement-related theta over immobility-related theta are consistent with reported differences in rabbits (Whishaw, 1976). Our findings are also consistent with previous suggestions that drugs acting on GABAergic receptors tend to modify the slope of the RPO-stimulation intensity versus HPC theta frequency function (McNaughton et al., 2007), and that all anxiolytics tested so far in the locomotion speed paradigm modify the y-intercept of the locomotion speed versus HPC (and mEC) theta frequency function (Wells et al., 2013, Korotkova et al., 2018). However, we also observed a non-specific large increase in the y-intercept of the locomotion speed versus HPC theta frequency relationship, which appears to be related to repeated testing on the same day. This effect has been reported previously in the mEC and the HPC at differing magnitudes (Newman et al., 2013, Wells et al., 2013, Monaghan et al., 2017). Baseline cholinergic-dependent theta frequency has been theorised to serve as the y-intercept of the speed vs frequency function (Burgess, 2008, Korotkova et al., 2018). However, scopolamine (0.5 mg/kg) marginally increases the y-intercept but significantly flattens the slope of the locomotion speed versus mEC theta frequency function (Newman et al., 2013), while selective genetic ablation of medial septal cholinergic neurons does not change the y-intercept despite increasing the slope in the HPC (Magno et al., 2017). In addition, scopolamine (up to 0.9 mg/kg) did not have a significant effect on RPO-elicited HPC theta frequency (McNaughton and Sedgwick, 1978). Therefore, it is unlikely that the y-intercept of the locomotion speed vs theta frequency function can be accounted for by cholinergic-dependent theta mechanisms. At present, there is no single theoretical framework to account for differential changes in y-intercept and slope in RPO stimulation intensity or locomotion speed-related HPC/mEC theta frequency.

The neurobiological effects of EtOH are multi-faceted. How HPC and cortical theta oscillations are generated are incompletely understood. Data from the current experiment fills in gaps in our understanding how low doses of EtOH may impact on neural oscillations, how the changes may be related to behaviour and how they modify the theta generating circuitry in context of being an anxiolytic. Future studies would benefit from continuous, real-time monitoring of neural activities and intra-cerebral ethanol concentrations (Rocchitta et al., 2012) to better attribute direct actions of ethanol on the brain. Given the prevalence of anxiety and related mood disorders, further work on the nature of the double-dissociation of how two HPC theta eliciting paradigms are modified by anxiolytics (i.e. by slope or y-intercept modification) may provide additional insights to their mode of action and thus biological bases of symptoms they treat.

## Acknowledgements

Materials support was provided by the University of Otago. Neither of the authors have conflicts of interest to declare.

